# The effects of host availability and fitness on *Aedes albopictus* blood feeding patterns in New York

**DOI:** 10.1101/2021.05.13.443995

**Authors:** Kara Fikrig, Elisabeth Martin, Sharon Dang, Kimberly St Fleur, Henry Goldsmith, Sophia Qu, Hannah Rosenthal, Sylvie Pitcher, Laura C. Harrington

## Abstract

*Aedes albopictus* is a competent vector of numerous pathogens, representing a range of transmission cycles involving unique hosts. Despite the important status of this vector, variation in its feeding patterns is poorly understood. We examined the feeding patterns of *Ae. albopictus* utilizing resting collections in Long Island, New York, and contextualized blood meal sources with host availability measured by household interviews and camera traps. We identified 90 blood meals, including 29 human, 22 cat, 16 horse, 12 opossum, 5 dog, 2 goat, and 1 rabbit, rat, squirrel and raccoon. Our study is the first to quantitatively assess *Ae. albopictus* feeding patterns in the context of host availability of wild animals in addition to humans and domestic animals. Host feeding indices showed that cats and dogs were fed upon disproportionately often compared to humans. Forage ratios suggested a tendency to feed on cats and opossums and to avoid raccoons, squirrels, and birds. This feeding pattern was different from another published study from Baltimore, where *Ae. albopictus* fed more often on rats than humans. To understand if these differences were due to host availability or mosquito population variation, we compared the fitness of Long Island and Baltimore *Ae. albopictus* after feeding on rat and human blood. In addition, we examined fitness within the Long Island population after feeding on human, rat, cat, horse, and opossum blood. Together, our results do not show major mosquito fitness differences by blood hosts, suggesting that fitness benefits do not drive Northeastern *Ae. albopictus* feeding patterns.

## Introduction

*Aedes albopictus* is a globally invasive mosquito of human and veterinary health importance. This species is capable of transmitting over 20 pathogens in laboratory assays^1^, and is a confirmed natural vector of dengue, Zika, and chikungunya viruses, and dog heart worm^1, 2^. *Aedes albopictus* is a suspected vector of numerous additional viruses, including Eastern equine encephalitis and West Nile due to virus detection in field-collected mosquitoes, although there is no direct evidence of transmission to humans yet^1^. These pathogens encompass vastly different transmission cycles, including anthroponoses (e.g. Zika: human to mosquito) and zoonoses (e.g. West Nile virus: primarily bird to mosquito to human; dog heartworm: dog or wild canid to mosquito). In light of the broad vector potential of *Ae. albopictus* and variation in feeding patterns in nature, it is critical to perform host feeding studies in locations relevant to human and animal health risk.

Variation in mosquito host feeding patterns can be influenced by a number of factors including innate host preference, environmental conditions, host availability, and the design of the studies themselves. These factors may explain the variation in host feeding reported for *Ae. albopictus* in the literature.

Published results for *Ae. albopictus* range from generalist or mammalophilic to highly anthropophagic (=human feeding) feeding patterns. For example, a high percentage of mosquitoes with human-derived blood meals were identified in tropical countries such as Thailand (100%) and Cameroon (99.4%)^3, 4^. In Thailand, aspirator collections were conducted around human dwellings, however, in Cameroon, mosquitoes were collected at a leisure and equestrian center, both of which were surrounded by human dwellings. In some parts of the USA, human feeding frequency was much lower, such as at a tire dump in Missouri (6.5%), urban Baltimore, Maryland (13.6%), urban and rural sites in Hawaii (18.1%), and suburban North Carolina (20%)^5, 6, 7, 8^. Additional studies have reported moderate human feeding rates such as in urban and peripheral sites in Brazil, urban and suburban Japan, and suburban New Jersey, USA^9, 10, 11^. Of those populations that did not feed predominantly on humans, most fed on a diverse array of animals, with the exception of Baltimore, where a striking number of *Ae. albopictus* fed on rats (72.3%)^6^.

One notable consistency amongst all published studies (with a sample size over 75) is a tendency for *Ae. albopictus* to feed primarily on mammals compared to birds and reptiles^3, 4, 5, 6, 7, 8, 10, 11, 12, 13, 14, 15, 16, 17, 18^. About half of studies report feeding on birds at low rates (1.7% to 25.6% of all blood meals)^5, 7, 8, 11, 13, 14, 16, 17, 18^. A tendency to feed even sporadically on birds is particularly important because of their role as amplifying hosts of arboviruses such as West Nile and Eastern equine encephalitis.

Host availability is rarely considered in the design of mosquito blood feeding studies despite its importance in driving mosquito blood feeding patterns and thus interpreting study results. In Italy, *Ae. albopictus* from urban and rural sites had replicable differences in feeding patterns, mirroring differences in host availability at these sites^14^. However, the authors only made qualitative note of site type and did not quantify host availability. We are aware of only two published studies (in North Carolina and Brazil) that have quantitatively assessed the link between host availability and blood feeding for *Ae. albopictus*^8, 13^. Their results do not provide a clear picture of whether *Ae. albopictus* feeds disproportionately often on humans compared to other mammals, with results varying depending on measurement type, stratification level, and non-human animal in question.

In addition to host availability, host attraction may be a major driver influencing blood feeding patterns^19^. Unfortunately, only two published studies have explored host attraction in *Ae. albopictus*^20, 21^. The authors reported higher attraction to humans compared to numerous other species including dogs and chickens. Preferential attraction to hosts is determined genetically, and may evolve as a result of elevated mosquito fitness after ingesting a given species’ blood^19, 22^. This has been demonstrated for *Ae. aegypti*, which maximizes reproductive fitness on human blood, its preferred host^23^. Only two studies have addressed the impact of blood from different species on *Ae. albopictus* egg production^24, 25^, but none have compared both survival and fecundity using the most ecologically relevant hosts.

We sought to determine *Ae. albopictus* feeding patterns in suburban and farm landscapes along its front of active northward expansion in New York State^26^. Our aim was to investigate these feeding patterns in the context of host availability and their consequences for mosquito fitness. Ultimately, we wanted to fill a gap in our understanding of *Ae. albopictus* feeding ecology along its Northeast USA range limit and how it might relate to public health risk. To rigorously address blood feeding patterns, we performed host censes to calculate host feeding indices and forage ratios. We then assessed whether fitness of Long Island, NY *Ae. albopictus* varied by host blood species ingested in the laboratory through a series of life table studies. To explore population differences, we compared fitness of Long Island and Baltimore populations fed human and rat blood meals.

## Methods

### Field Sites

Eight sites were selected in Suffolk County on Long Island, NY: four farms and four residential areas, each containing between nine and seventeen collection properties. *Ae. albopictus* has been present in Suffolk County since 2004, although its distribution is not uniform or complete across the county (Moses Cucura, pers comm). Residential sites were selected based on *Ae. albopictus* presence reported by the Suffolk County Vector Control and Arthropod-Borne Disease Laboratory and larval distribution data^27^. All residential sites were suburban, with variable human population density: Central Islip (1,853 people/sq km), Bay Shore (1,853 people/sq km), Babylon (1,660 people/sq km), and Hauppauge (734 people/sq km). All four farms were partially bordered by suburban residential and forested natural landscapes.

### Collection

Weekly collections were conducted at each site between 20 June and 15 August, 2018 with large custom-designed aspirators (30.5 cm diameter, 114 cm height, 12 V PM DC 2350 RPM, 1/35 Horse power, 3.7 amp motor)^3^. One mosquito was collected with a hand net while host-seeking near collectors and was included in analysis because blood was partially digested upon collection. Mosquitoes were immobilized in acetone-treated jars (3 min) and sorted in the field to remove non-mosquito by-catch. The samples were transported on ice to the laboratory for identification according to a taxonomic key^28^. *Aedes albopictus* were considered engorged if blood was visible in the abdomen upon examination. Mosquitoes were stored at −20°C and transported to Cornell University on dry ice for blood meal identification.

### Blood Meal Identification

Abdomens were removed from mosquitoes using forceps and transferred to sterile microcentrifuge tubes. To avoid cross-contamination, forceps were dipped in ethanol and flame-sterilized between each sample. DNA was extracted from abdomens using Qiagen Puregene Cell kit (Qiagen Sciences, Germantown, MD, USA). To identify blood meals, we amplified templates from the vertebrate-specific cytochrome c oxidase subunit I (*COI)* “barcoding” gene. Primers designed by Reeves et al. (2018) were used to amplify a 395 base pair amplicon^29^(Table 1).

**Table 1:** Primer sequences designed by Reeves et al. (2018)

| Primer Name | Sequence |
| --- | --- |
| VertCOI_7194_F | 5'- CGM ATR AAY AAY ATR AGC TTC TGA Y -3' |
| Mod_RepCOI_R | 5'- TTC DGG RTG NCC RAA RAA TCA -3' |

Other Reeves *COI* primers were not used due to co-amplification of *Ae. albopictus* DNA. Co-amplification is a recurrent issue with identifying *Ae. albopictus* blood meals due to matching sequences between its own genome and primers designed for use in blood meal studies of other mosquito species^15^. Notably, cytochrome b primers designed by Egizi et al. (2013) were used initially, but due to low success rate in our hands, we switched to the Reeves primers^29^. Three blood meals identified with the Egizi primers were not successfully amplified by the Reeves primers; results with both primer sets were combined for our data analysis.

PCR conditions were slightly modified from Reeves et al. (2018) in order to minimize co-amplification of *Ae. albopictus* DNA and maximize amplification of desired amplicon^29^. Reactions were performed with total volume of 20 μL, consisting of 10 μL of 2.0X Apex Taq RED Master Mix (Genesee Scientific Corp., San Diego, CA), 0.75 μL of VertCOI_7194_F forward primer (10 μM), 0.75 μL of Mod_RepCOI_R reverse primer (10 μM), 6.5 μL sterile nuclease-free H_2_O, and 2 μL of extracted DNA. Most reactions were conducted with the following thermocycling conditions: 94°C for 3 min, followed by 40 cycles of 94°C for 40 s, 53.5°C for 30 s, and 72°C for 60 s, and a final extension step at 72°C for 7 min. The annealing temperature was modified from Reeves et al. (2018) in order to minimize amplification of *Ae. albopictus* DNA according to a temperature gradient test conducted on positive (human-fed) and negative (non-fed) mosquito controls. Conditions were further modified for a subset of reactions to optimize amplification: 94°C for 3 min, followed by 5 cycles of 94°C for 40 s, 45°C for 30 s, and 72°C for 60 s, and then 35 cycles of 94°C for 40 s, 48.5°C for 30 s, and 72°C for 60 s, and a final extension step at 72°C for 7 min. All reactions were conducted alongside a positive (human-fed mosquito) and negative (sterile nuclease-free water) control. PCR products (5 μL) were loaded onto a 1% agarose gel stained with gelRED, electrophoresed, and visualized with UV light (Mighty Bright, Hoefer Scientific Instruments, San Francisco, CA, USA).

Samples with positive bands after gel electrophoresis were purified with FastAP and Exonuclease (ThermoFisher Scientific, Waltham, MA, USA) and submitted for Sanger sequencing at the Cornell University Biotechnology Resources Center. Sequences were compared to the available database in NCBI Basic Local Alignment Search Tool (BLASTn) and were identified to a source if matches were ≥98% with a sequence of known origin (with the exception of an eastern gray squirrel (*Sciurus carolinensis*) sequence, which had a 95.5% match).

### Host Availability

#### Household Interviews

To estimate host availability, household interviews were conducted weekly at time of collection (see Supplemental Materials S1). Residents were asked about the number of people and pets living in their house and the amount of time spent outside by species that day and the two days prior. Interviews were conducted in English or Spanish depending on homeowner preference.

#### Camera Traps

Two motion-triggered camera traps (Moultrie M-880, #MCG-12691, Calera, AL, USA) were set at each site from 16 July to 13 August 2018 on selected properties in residential sites and different locations within farm sites. Cameras were operated according to the setting, height, and angle specifications described by Linske et al.^30^, with the exclusion of scent lures. Camera data were used to estimate host abundance by determining the number of animal encounters with the camera per trap day. If a given species was photographed within 30 min of the last image of that animal, it was considered the same individual and was not counted separately. If multiple individuals were captured in one image within 30 min of last sighting, the count was equal to the maximum number captured together in an image.

#### Fitness by Host Species

##### Mosquito Rearing

Mosquitoes were collected from four towns on Long Island, NY and reared in colony for six to ten generations. Eggs from Baltimore, MD (between F_3_ and F_6_ depending on replicate) were reared synchronously with the Long Island colony in order to assess between population differences. For each replicate, eggs were vacuum hatched, provided with a pinch of pulverized fish food (crushed Cichlid Gold™ fish food pellets; Hikari, Himeji, Japan), and one day later, separated into trays of 200 larvae, with 1L of distilled water, and 4 Cichlid Gold™ fish food pellets. Adult mosquitoes were maintained in an environmental chamber (28°C, 71.9% ± 9.5% relative humidity, 10 hr light, 10 hr dark, 2 hr dusk/dawn). Cups of 200 pupae were placed into cages inside the chamber, and upon eclosion, 10% sucrose was provided for 2-4 d. Males were removed and sucrose was replaced with distilled water for 1 d prior to blood feeding.

##### Blood

Human (Lampire Biologicals; Pipersville, PA, USA), opossum (The Janet L. Swanson Wildlife Health Center; Ithaca, NY, USA), rat (The Center for Animal Resources and Education, Cornell University), cat (The Center for Animal Resources and Education at Cornell University; Ithaca, NY, USA) and horse (Lampire Biologicals; Pipersville, PA, USA) blood treated with anticoagulant (sodium citrate) was stored at −20°C upon arrival. Blood was thawed in warm water immediately before use. Mosquito blood feeding was conducted with artificial feeders (water reservoir at 37°C and de-salted sausage casings as membrane) as described previously^31^.

#### Within-population differences of Long Island Ae. albopictus

In order to determine whether fitness advantages for different host blood sources influence feeding patterns of Long Island *Ae. albopictus*, we assessed fecundity and survival of females after feeding on human, cat, horse, opossum, and rat blood. These species were chosen based on commonly identified blood sources in our study or in Baltimore, MD^6^.

##### Fecundity and Survival

Fully engorged mosquitoes (approximately 35 per blood species per replicate and 3-4 replicates per group) were gently transferred individually into 0.5L paper cups with a dry oviposition vessel. Mosquitoes were maintained in the environmental chamber as described above. One day after blood feeding, strained larval rearing water was added to oviposition vessels to encourage egg lay. No additional water or sugar was provided. Each mosquito was checked daily for presence of eggs (first day of egg lay) and mortality until all females had died. Total number of eggs laid per female was recorded at the end of experiment. Dead mosquitoes were frozen at −20°C and later dissected to determine number of mature retained eggs, if any. We compared the total eggs produced (retained + laid eggs). In replicate two, mosquitoes with a large number of retained eggs were not counted and were therefore not included in the egg analyses but were included in survival analyses. For individuals where egg retention data was not available, number of eggs laid was used. The following blood types were tested: replicate one included human, rat, cat, and horse; replicates two and three included human, rat, cat, horse, and opossum; replicate four included human, rat and opossum.

#### Between-population differences of Long Island and Baltimore Ae. albopictus

Because of the striking differences in field-collected host blood meal sources between our study and a prior Baltimore study^6^, we assessed whether fitness varied between *Ae. albopictus* from these two locations after feeding on rat (source of 72.3% of blood meals in Baltimore, and 1.1% of blood meals in our current study of mosquitoes on Long Island) and human blood (source of 13.6% of blood meals in Baltimore and 32.2% in Long Island)^6^.

##### Fecundity and survival

Long Island and Baltimore *Ae. albopictus* were fed rat and human blood and observed synchronously. The rat and human-fed Long Island individual mosquitoes from replicates 1-3 of the within-population fitness assessment described above were used to compare both between-population fitness of Long Island and Baltimore *Ae. albopictus* and within-population fitness of Long Island *Ae. albopictus*. The wing length of a subset of Long Island and Baltimore individuals was measured to control for body size differences between the two colonies^32, 33^.

### Data Analysis

#### Host availability

##### Residential Host Feeding Index

Abundance and time-weighted host feeding indices (HFI) were calculated using blood meal identification data from residential areas and household interview data for humans, cats and dogs. Feeding indices were calculated according to equations described by Kay et al. (1979) and modified by Richards et al. (2006) as follows^8, 34^:

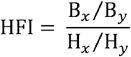

where B_*x*_ and B_*y*_ represent the average number of blood meals from host *x* and host *y* per household and H_*x*_ and H_*y*_ represent the average number of host *x* and host *y* residing per household. Averages were calculated with data from households positive for at least one bloodmeal. Data were aggregated across all four residential sites because household and site-specific calculations frequently resulted in non-real values due to zeroes in the denominators.

A time-weighted feeding index ^8^ was calculated as follows:

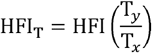

where T_*y*_ and T_*x*_ represent the time spent outside by hosts *y* and *x*, respectively. When household interview data was missing on the date of bloodmeal collection (26 of 66 surveys), the average of all other interview responses from that household was used as an approximation.

An HFI or HFI_T_ greater than 1 indicated that host *x* was fed upon more often than expected compared to host *y* given their abundance or time spent outside. An HFI or HFI_T_ equal to 1 indicated that the hosts were fed upon in proportion to their availability and an HFI or HFI_T_ less than 1 indicated that host *y* was fed upon more often than expected compared to host *x*. Note that while an HFI or HFI_T_ greater or less than 1 may reflect *Ae. albopictus* preference, it does not conclusively demonstrate it, as we cannot rule out influences from other factors such as host defenses, timing of host availability, or host location in the yard.

##### Residential Forage Ratio

Forage ratios are another method for determining host feeding frequency by host availability^3^. In our study, these were calculated using blood meal identification data and camera trap images from residential sites. Forage ratios were calculated for each animal species that was captured by camera traps as follows^35^:

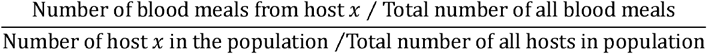

In the case of this study, the proportion of all hosts represented by host *x* was approximated by the proportion of all camera trap images that were taken of host *x*. Because camera traps were only placed in 2 properties per site, forage ratio calculations were limited to animals that tend to cross freely between yards, including all wild animals and cats, but excluding humans and dogs.

A forage ratio greater than one suggests that the host was fed upon more often than expected given its abundance and less than one suggests that the host was fed upon less often than expected. A forage ratio equal to one indicates that the host was fed upon in proportion to its abundance in the population. As with host feeding indices, forage ratios may reflect preference but do not prove it because the same sources of bias may impact these results.

##### Farm Host Availability

At the farm sites, host feeding indices and forage ratios were not calculated due to small sample sizes and technical difficulties of defining host availability. Interviews of human and domestic animal availability were only conducted once at farms during the last week of collections. Farm owners could not accurately estimate human exposure due to unpredictable influx of people on site for riding lessons and farm work. Animal exposure could not be reliably measured because of inconsistent use of fenced paddocks and semi-enclosed barns. Camera traps were positioned in order to picture wild animals at the outskirts of the fenced paddocks and therefore did not often picture domestic farm animals. Interview and camera trap data is reported for each but are only qualitatively compared to blood meal data; no further calculations were conducted.

#### Life table studies-fitness by host species

##### Within-population differences

The effect of host blood source on egg production (fecundity) was assessed with a linear model, including replicate and mosquito survival as covariates. The effect of host blood source on mosquito survival was also determined using a linear model, including replicate as a covariate. Estimated marginal means *post hoc* analyses were conducted using the emmeans package^36^. Survival curves were created with the average proportion surviving across the replicates and compared for each host blood species. The basic reproductive rate (R_0_) was calculated for each blood type and replicate according to previously described equations ^37^. The effect of blood type on R_0_ was compared via a linear model.

##### Between-population differences

Egg production and survival were compared between human/rat, Long Island/ Baltimore groups using linear models, as described above. However, in this case, number of eggs produced by each individual was divided by average wing length of the cohort, reported as eggs per mm wing length (eggs/mm wl), in order to control for the effect of body size, which differed between Baltimore and Long Island colonies despite identical rearing.

### Ethics approval

Survey protocols were reviewed and considered exempt by Cornell University’s Institutional Review Board (IRB).

## Results

### Blood Meal Identification

3,241 *Ae. albopictus* were collected over the course of the summer (1,575 female and 1,666 male) and 182 (14% of aspirator-collected females) were blood-fed. Of these, 152 blood meals were less than half digested. Host identity was successfully assigned to 90 samples (49.5%), including 29 human (*Homo sapiens;* 32.2%), 22 cat (*Felis catus;* 24.4%), 16 horse (*Equus caballus;* 17.8%), 12 opossum (*Didelphis virginiana;* 13.3%), 5 dog (*Canis lupus familiaris;* 5.6%), 2 goat (*Capra hircus;* 2.2%), and 1 each of rabbit (*Sylvilagus floridanus*; 1.1%), rat (*Rattus norvegicus*; 1.1%), squirrel (*Sciurus carolinensis*; 1.1%), and racoon (*Procyon lotor*; 1.1%). When divided into residential (n=66) and farm sites (n=24), most of the residential blood meals were from humans (40.9%), followed by cat (31.8%) and opossum (18.2%). The majority of farm blood meals were from horses (66.7%), followed by human (8.3%) and goat (8.3%) (Figure 1).

**Figure 1.**
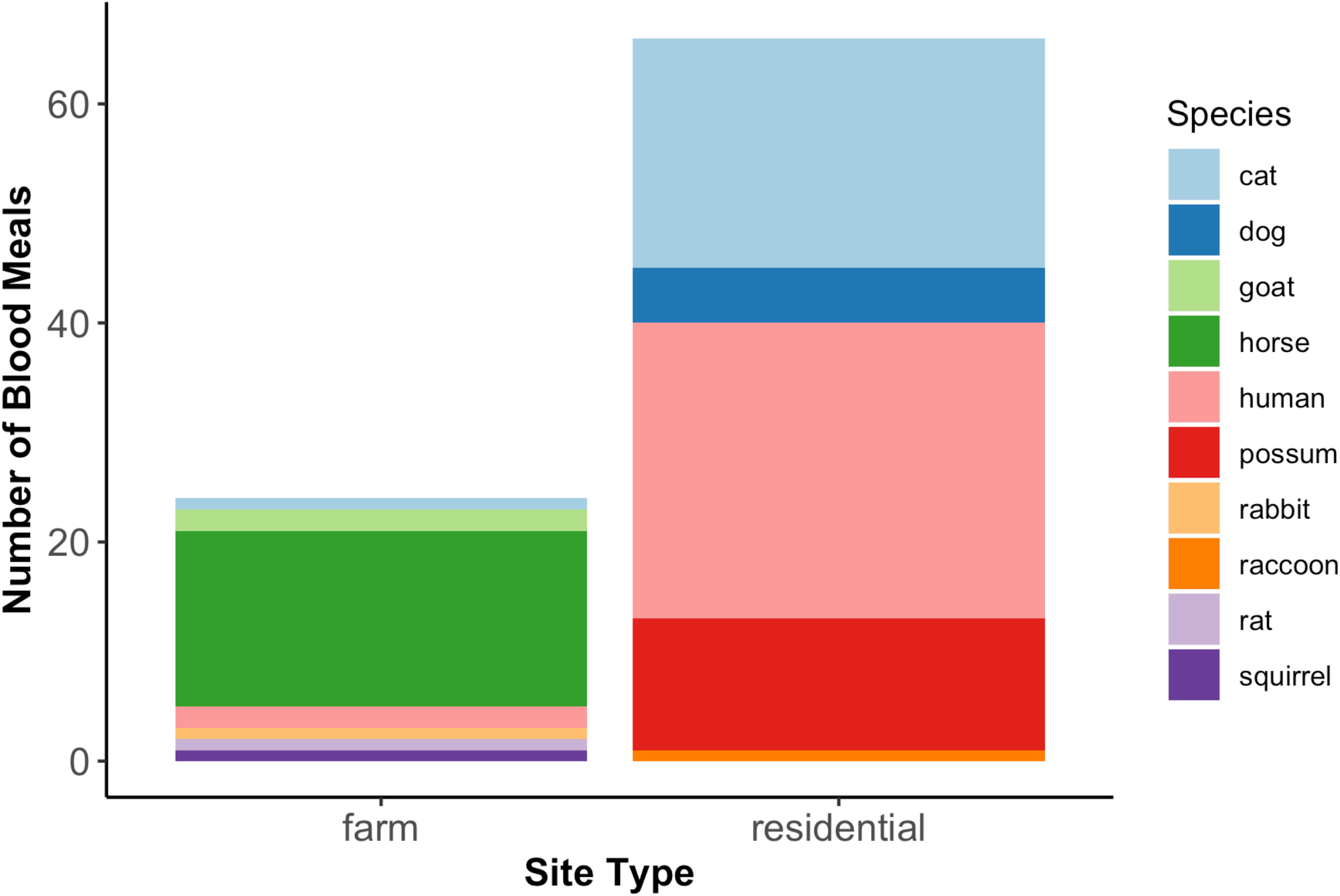
Distribution of blood meals by site type. At residential sites, most *Ae. albopictus* fed on human, followed by cat and opossum. At farm sites, the majority fed on horse, followed by human and goat.

### Host availability

#### Residential Host Feeding Index

Household interview and blood meal data were used to calculate host feeding indices (HFIs), indicative of relative tendency to feed on certain vertebrate hosts at all residential properties where blood meals were collected (n=28) (Table 2). The most human blood meals were collected per property (0.96 ± 0.21), followed by cat (0.75 ± 0.17), and dog (0.18 ± 0.09). Similarly, there were the most human residents per property (3.18 ± 0.36), followed by cat (0.39 ± 0.19), and dog (0.29 ± 0.10). However, cats spent the most time outside over the 2 days prior to collection (278.74 ± 232.93 min), followed by humans (234.26 ± 49.83 min), and dogs (53.61 ± 22.05 min). The standard error in cat time was large because some individuals were outdoor cats (24 hrs/d) while others were only allowed outside for short periods of time.

**Table 2.** Mean (±SE) number of blood meals, residents, and time spent outside for humans, cats and dogs per property

| Mean (± SE) per property | | | |
| --- | --- | --- | --- |
| Host | Blood meal | Residents | Time outside (min) |
| Human | 0.96 (0.21) | 3.18 (0.36) | 234.26 (49.83) |
| Cat | 0.75 (0.17) | 0.39 (0.19) | 278.74 (232.93) |
| Dog | 0.18 (0.09) | 0.29 (0.10) | 53.61 (22.05) |

Mean numbers of blood meals and residents were used to calculate pairwise comparisons of feeding between humans, cats, and dogs through abundance and time-weighted HFIs (Table 3). Human vs cat HFI and HFI_T_ both demonstrate a tendency to feed on cats compared to humans (0.16 and 0.20). Likewise, human vs dog HFI and HFI_T_ both suggest that *Ae. albopictus* feeds disproportionately often on dogs compared to humans (0.49 and 0.14). However, cat vs dog HFI and HFI_T_ produced opposite results: according to abundance measures, cats were fed upon disproportionately more often compared to dogs (3.05), but when time-weighted, dogs were fed upon disproportionately more often compared to cats (0.73). On average, cats spent much more time outside than dogs, causing the directionality change of the index. Furthermore, neither HFI metric demonstrates a particularly strong deviance from the expected feeding proportions, suggesting that *Ae. albopictus* may not have a strong preference between cats and dogs.

**Table 3.** Abundance and time-weighted host feeding indices

Abundance and time-weighted host feeding indices
| Index | Human vs Cat | Human vs Dog | Cat vs Dog |
| --- | --- | --- | --- |
| HFI | 0.16 | 0.49 | 3.05 |
| HFI<sub>T</sub> | 0.20 | 0.14 | 0.73 |

#### Residential Forage Ratio

Forage ratios (FRs) were calculated from camera trap data at the 4 residential sites for all animals for which camera trap images were taken or blood meals collected (Table 4). Cats and opossums were fed upon more often than expected given their relative abundance in the host population. Of all residential blood meals taken from free roaming species (i.e., not humans and dogs), 65.7 ± 10.2% were derived from cats, but only 27.4 ± 10.9% of all images were taken of cats, resulting in a 3.56 ± 0.98 FR (above the FR=1 threshold to infer preference). Opossum blood meals accounted for 31.8 ± 10.8% of all blood meals but no opossums were pictured, resulting in an undefined FR, suggesting preference for opossums. Raccoons, the other nocturnal animal, were pictured often (24.8 ± 16.4% of all images) but only represented 2.5 ± 2.5% of all blood meals, resulting in a FR below 1 (0.046 ± 0.046), suggesting avoidance. Squirrels and birds were also pictured often (21.6 ± 10.5% and 26.2 ± 11.2% respectively) but no blood meals were collected at residential sites, resulting in a FR of 0, suggesting avoidance.

**Table 4.** Mean (± SE) percentage of all blood meals, percentage of all animals and forage ratio for all animals for which camera trap images were taken or blood meals collected at residential sites (n=4) in Suffolk Country, NY.

| | Mean (± SE) | | |
| --- | --- | --- | --- |
|  | % of all blood meals | % of all images | Forage Ratio |
| Cat | 65.7 (10.2) | 27.3 (10.9) | 3.6 (1.0) |
| Possum | 31.8 (10.8) | 0 (0) | $\infty^*$ |
| Raccoon | 2.5 (2.5) | 24.8 (16.4) | 0.05 (0.05) |
| Squirrel | 0 (0) | 21.6 (10.5) | 0 (0) |
| Bird | 0 (0) | 26.2 (11.2) | 0 (0) |

\*FR was infinite because division by zero is undefined

#### Farm Host Availability

Approximate numbers and time spent outside for humans and domestic animals were reported by the farm owners. At Farm A, approximately nine people spent time at the farm for a total of 52 hours per day. The farm also had 40 horses, spending a total of 70 hrs/d outside. At Farm A, 3.6% of camera trap images were of cats, 67.9% of raccoons, 17.9% of foxes, 3.6% of deer, and 7.1% of squirrels. Blood meals collected at Farm A included 6 horse and 1 squirrel.

Farm B estimated that 30 people (180 hrs), 100 horses (200 hrs), 2 dogs (26 hrs), and 2 goats (26 hrs) were outside on the property per day. Of all camera trap images at Farm B, 37.1% were of cats, 44.3% of raccoons, 4.1% of opossums, 5.2% of deer, 5.2% of squirrels, and 4.1% of rabbits. The blood meals consisted of 5 horses, 1 human, and 1 rabbit.

Farm C estimated that 7 people (11 hrs), 46 horses (420 hrs), 2 dogs (12 hrs), 18 chickens (171 hrs), 4 ducks (38 hrs), and 1 goose (24 hrs) spent time outside per day. The most images were taken of cats (48.8%), followed by birds (23.3%), raccoons (14.0%), squirrels (9.3%) and rabbits (4.7%). Blood meals included 4 horses and 1 cat.

Farm D estimated that 3 people (14 hrs), 8 horses (48 hrs), 2 dogs (8 hrs), 20 goats (260 hrs), 4 sheep (52 hrs), 1 alpaca (24 hrs), 1 llama (24 hrs), 20 rabbits (260 hrs), 9 ducks (117 hrs), and 30 chickens (720 hrs) spent time outside per day. The camera trap pictured raccoons (33.3%) and birds (66.7%). Blood meals collected included: 2 goat, 1 horse, 1 human, and 1 rat.

Despite the diversity of hosts available at the 4 farm sites, the predominant blood meal identified at three of these sites was horse. The fourth farm was an anomaly, with more blood meals collected from goats than horses, but it was also the only farm where more goats were available than horses. Once again, raccoons were pictured at all sites, but no blood meals were collected, further suggesting avoidance of this animal. Birds were pictured frequently at 2 sites, and no blood meals collected, also suggesting avoidance.

### Fitness by Host Species

#### *Within-population differences of Long Island* Aedes albopictus

The proportions of *Ae. albopictus* that laid and retained mature eggs and mean (± SE) number of eggs produced are reported in Table 5.

**Table 5:** Egg production by blood meal source for Long Island *Ae. albopictus*

| **Blood Source** | **Proportion which laid eggs (%)** | **Proportion with retained eggs (%)\*** | **Mean eggs produced (± SE)** |
| --- | --- | --- | --- |
| Cat | 57/89 (64.0) | 10/89 (11.2) | 40.3 (4.0) |
| Horse | 70/97 (72.2) | 11/97 (11.3) | 48.5 (3.9) |
| Human | 104/121 (86.0) | 23/121 (19.0) | 61.0 (2.9) |
| Rat | 100/122 (82.0) | 16/122 (13.1) | 53.5 (3.7) |
| Opossum | 64/86 (74.4) | 10/86 (11.6) | 58.7 (4.8) |

\*Includes mosquitoes with any number of retained eggs

Females that ingested cat blood resulted in lower fecundity compared to those fed human and opossum blood (β= −17.3, SE=5.3, *P*=0.01 and β= −20.9, SE=5.9, *P*=0.004, respectively). There was no significant difference between any other blood group (Figure 2a). There was also no significant effect of survival time on number of eggs produced (although only one blood meal was provided in this study, which may limit impact of extended survival). On average, all blood groups began laying on day 3 post-blood meal, and all blood groups survived for 7-9 days. Notably, there were significant differences between replicates; mosquitoes in replicate 1 produced more eggs than replicate 2 and 3 (β= 30.67, SE=447, *P*<0.0001 and β= 35.69, SE=4.41, *P*<0.0001, respectively) and mosquitoes in replicates 2 and 3 produced fewer eggs than replicate 4 (β= −30.37, SE=5.37, *P*<0.0001 and β= −35.39, SE=5.38, *P*<0.0001, respectively).

**Figure 2.**
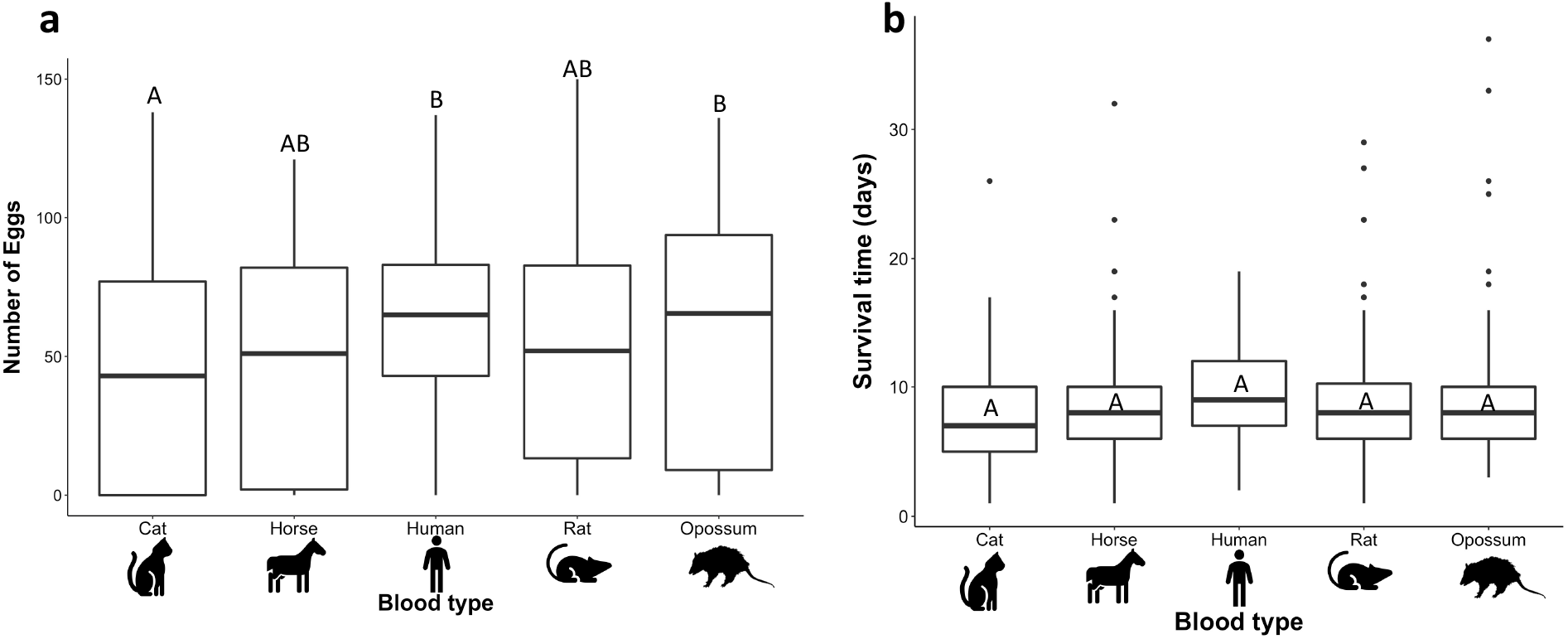
**a)** Box plot of number of eggs produced by Long Island mosquitoes fed cat, horse, human, rat, and opossum blood. Blood types that do not share a letter above boxes are significantly different. **b)** Box plot of survival time in days. Blood types that do not share a letter inside boxes are significantly different.

There were no significant differences in survival time between any of the host blood groups (Figure 2b). Mosquitoes fed cat blood survived (± SE) 7.6 (±0.45) days, horse-fed survived 8.6 (±0.48) days, human-fed survived 9.6 (±0.3) days, rat-fed survived 8.7 (±0.4) days, and opossum-fed survived 9.5 (±0.6) days. There were significant differences in survival by replicate: replicate 1 had higher survival than replicate 3 (β=1.7, SE=0.5, *P*=0.006) and replicates 1, 2, and 3 had lower survival than replicate 4 (β=-3.5, SE=0.6, *P*<0.0001; β= −4.5, SE=0.6, *P*<0.0001; β=-5.1, SE=0.6, *P*<0.0001 respectively). Daily survival curves averaged over the three or four replicates are presented in Figure 3.

**Figure 3:**
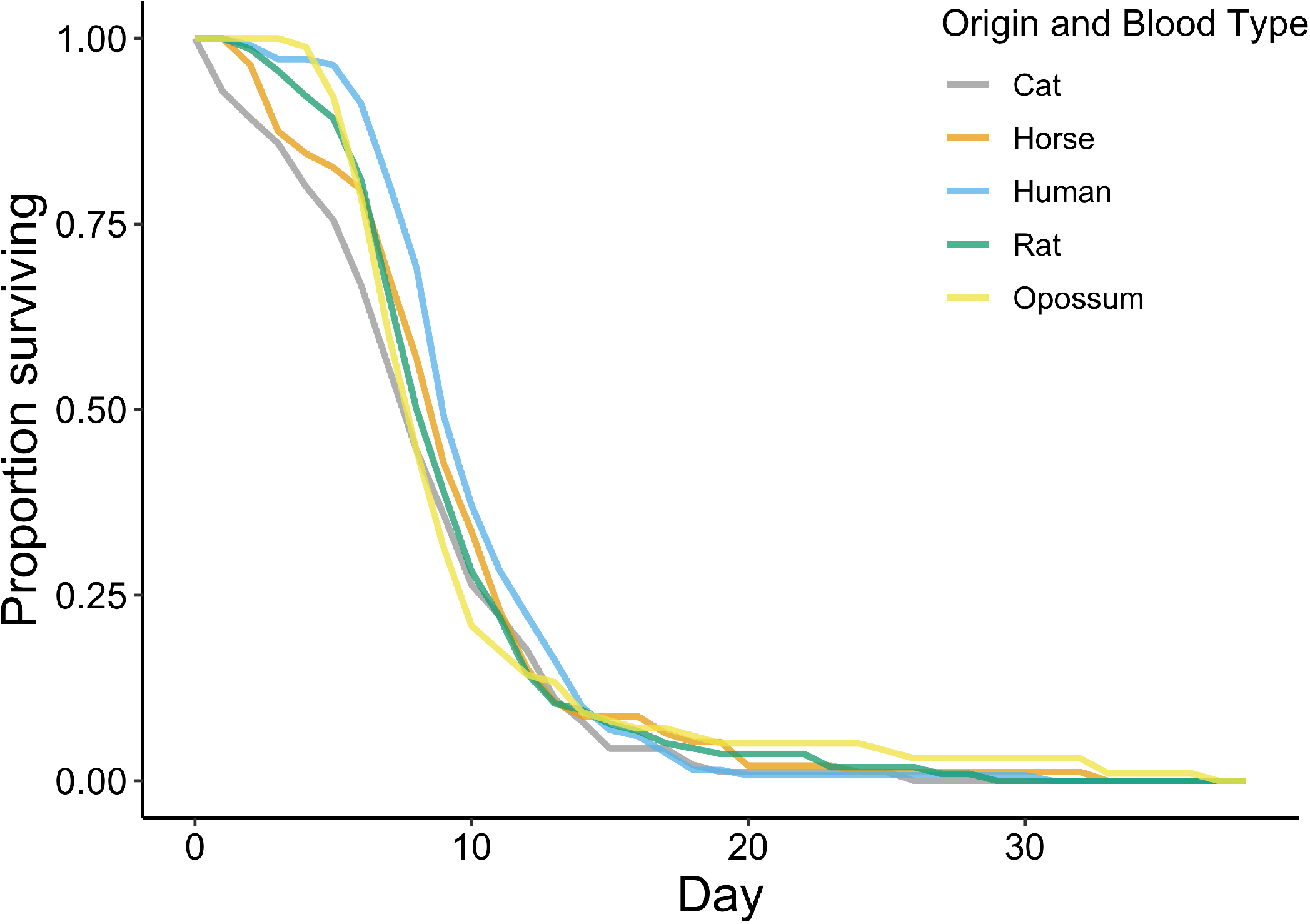
Survival of Long Island *Ae. albopictus* by host blood ingested

The mean (±SE) R_0_ across replicates was 19.5 (±6.5) for Long Island *Ae. albopictus* fed cat blood, 22.9 (±5.7) for horse blood, 29.7 (±4.1) for human, 27.1 (±8.9) for opossum, and 27.0 (±4.1) for rat. No significant differences in (R_0_) were found by host blood group.

#### Between-population differences of Long Island and Baltimore Ae. albopictus

The proportions of *Ae. albopictus* that laid and retained mature eggs, mean (± SE) eggs, and mean (± SE) eggs/mm wl is reported in Table 6.

**Table 6:** Egg production for Long Island and Baltimore *Aedes albopictus* females fed human or rat blood

| Origin and Blood Source | Proportion which laid eggs (%) | Proportion with retained eggs (%) | Mean eggs produced (± SE) | Mean eggs/mm w produced (± SE) |
| --- | --- | --- | --- | --- |
| Baltimore Human | 73/89 (82.0) | 12/89 (13.5) | 41.4 (3.2) | 14.7 (1.1) |
| Baltimore Rat | 70/95 (73.7) | 11/95 (11.6) | 38.2 (3.5) | 13.6 (1.3) |
| Long Island Human | 76/89 (85.4) | 17/89 (19.1) | 58.8 (3.6) | 20.7 (1.3) |
| Long Island Rat | 75/95 (78.9) | 13/95 (13.7) | 46.1 (4.0) | 16.2 (1.4) |

The only significant differences in eggs produced per mm wing length were between Long Island mosquitoes fed human blood and the three other groups (Figure 4a). Baltimore mosquitoes fed human (β= −6.0, SE=1.8, *P*=0.0008) and rat blood (β= −6.9, SE=1.8, *P*=0.0001) produced fewer eggs/mm wl than Long Island mosquitoes fed human blood. Long Island mosquitoes fed human blood produced more eggs per mm wl than those fed rat blood (β= 3.8, SE=1.8, *P*=0.03). Baltimore mosquitoes fed rat blood produced marginally fewer eggs/mm wl than Long Island mosquitoes fed rat blood (β= −3.1, SE=1.7, *P*=0.07). There was no significant difference in eggs produced/mm wl between Baltimore mosquitoes fed human and rat blood (β= 1.0, SE=1.8, *P*=0.6) or Baltimore mosquitoes fed human blood and Long Island mosquitoes fed rat blood (β= −2.1, SE=1.8, *P*=0.2).

**Figure 4.**
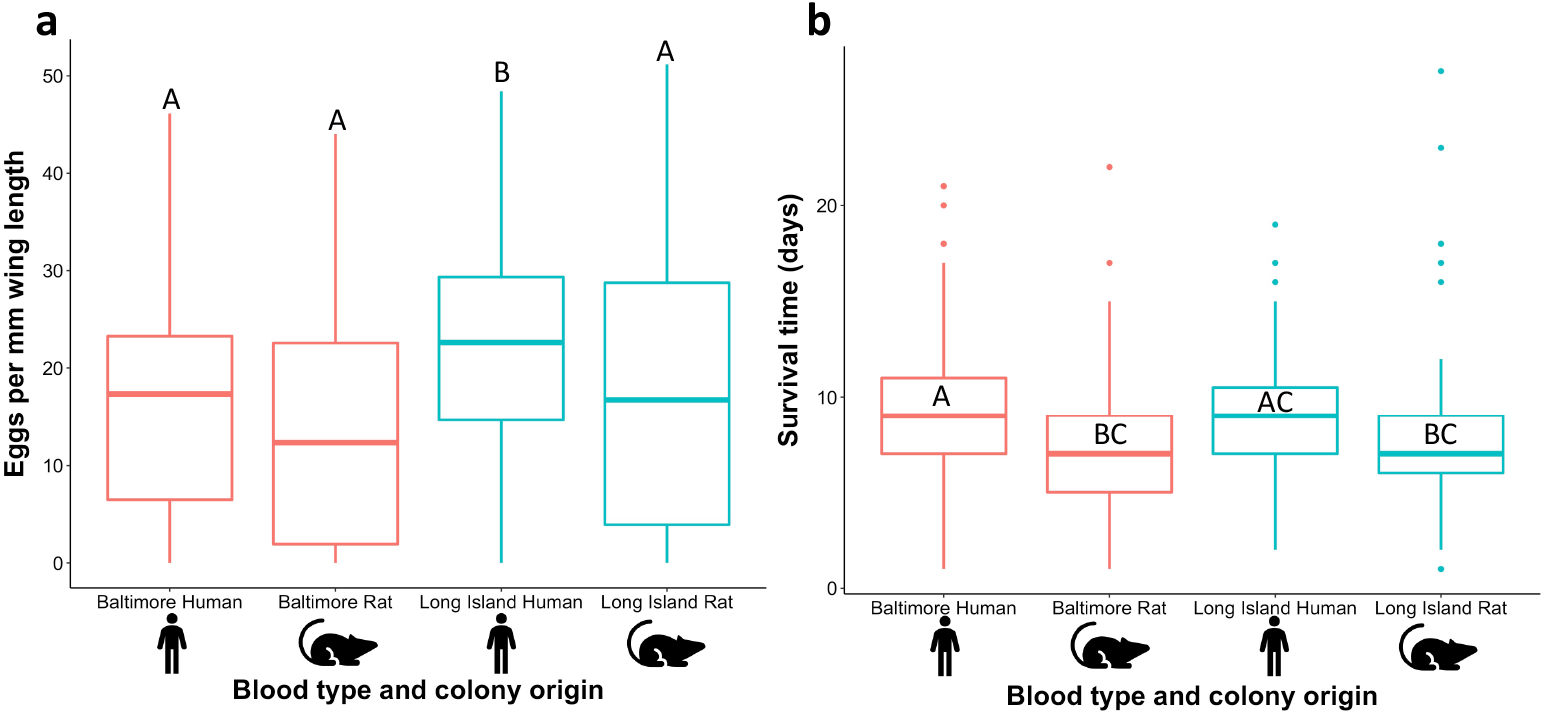
**a)** Box plot of number of eggs produced by Baltimore and Long Island mosquitoes fed rat and human blood. Groups that do not share a common letter are significantly different. **b)** Box plot of survival in days. Groups that do not share a common letter are significantly different.

The mean (± SE) survival time of Baltimore *Ae. albopictus* was significantly higher for human blood (9.6 days ±0.4) compared to rat blood (7.2 days ±0.4) (β= 2.3, SE=0.5, *P*=0.0001). The same survival trend was observed for Long Island *Ae. albopictus* where mosquitoes fed human blood survived marginally longer than those fed rat blood (9.0 days ±0.3 and 7.7 days ±0.4 respectively: β= 1.3, SE=0.5, *P*=0.08) (Figure 4b). Baltimore mosquitoes fed human blood survived significantly longer compared to Long Island mosquitoes fed rat blood (β=1.9, SE=0.5, *P*=0.002). Survival time was significantly lower for Baltimore mosquitoes fed rat blood compared to Long Island mosquitoes fed human blood (β=-1.7, SE=0.5, *P*=0.008). There was no significant difference in survival time between mosquitoes fed human blood from both sites (β= 0.6, SE=0.5, *P*=0.6) or fed rat blood from both sites (β= −0.4, SE=0.5, *P*=0.8). We did detect differences by replicate, where replicate 1 had a higher survival than replicate 2 (β= 1.4, SE=0.5, *P*=0.006) while paired replicates 1-3 and 2-3 had no significant difference in survival (β= 0.9, SE=0.4, *P*=0.1 and β= −0.6, SE=0.5, *P*=0.4 respectively). Daily survival curves averaged over the three replicates are presented in Figure 5.

**Figure 5:**
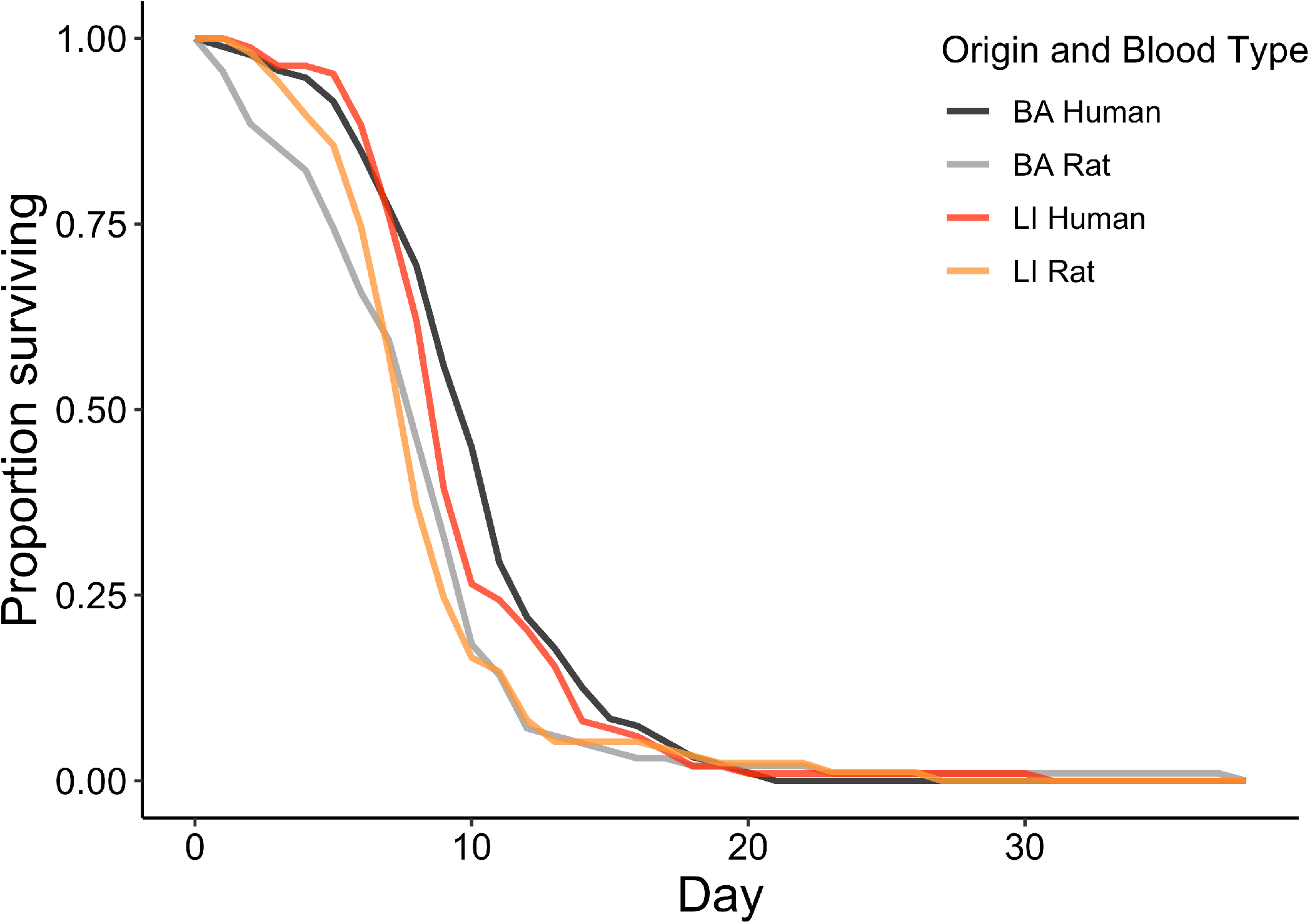
Survival of Baltimore and Long Island *Ae. albopictus* fed human and rat blood. Curves are averaged over three replicates.

The mean (±SE) basic reproductive rate (R_0_) (averaged over 3 replicates) of Baltimore *Ae. albopictus* fed human blood is 20.4 (±1.2), 19.7 (± 4.6) for Baltimore rat, 29.3 (± 5.7) for Long Island, and 24.5 (± 4.5) for Long Island rat. No significant differences occurred between R_0_ of any of the blood groups.

## Discussion

Mosquito feeding behavior plays a vital role in disease transmission, however, it can be difficult to understand and predict because there are diverse factors that influence feeding behavior in nature. We investigated the feeding patterns of the globally invasive vector, *Ae. albopictus*, in farm and residential habitats at the northern edge of its range in the United States. In tandem, we addressed two factors that may influence these patterns: host availability and variation in mosquito fitness from different host blood sources.

The ten host species we detected in *Ae. albopictus* blood meals from Long Island, NY are hosts previously reported for this species elsewhere in the world. The proportion of human blood meals (32.2%) identified in Long Island was lower than reported in many other locations worldwide^3, 4, 10, 11, 13, 14, 15, 16, 17, 18, 38^, but was higher than in other studies from the United States (Hawaii, Missouri, North Carolina, and Maryland)^5, 6, 7, 8^. More *Ae. albopictus* fed on cats in our study on Long Island than in any other location previously reported. The third most common host for this mosquito species on Long Island, the horse, has only been detected in four of sixteen previous *Ae. albopictus* blood meal studies and at lower levels^7, 8, 13, 14^. Similarly, the fourth most common host, opossum, has been reported in four previous studies, also at lower levels^5, 8, 10, 15^. Long Island *Aedes albopictus* fed less frequently on dogs compared to the representative proportion in numerous other studies^7^. Notably absent from the Long Island blood meals were cows, deer, and birds, all of which were present on at least one site in our study and have been detected in at least six previous blood meal studies. It is possible that a larger sampling of blood meals may have revealed these hosts, however, birds have also been absent from other studies in Northeastern USA^6, 10, 15^. Notably, only about half of collected blood meals were successfully identified to species, but the reason for the low success rate is unknown. It is possible that this may have biased the species that were identified, however, tests of primer versatility performed by Reeves et al. (2018) showed amplification for the majority of vertebrate species (90/93)^29^.

This is only the third study of *Ae. albopictus* blood feeding biology that quantitatively assessed host availability, and the first to do so with wild animals. Abundance and time-weighted host feeding indices (HFIs) calculated using household interview data revealed disproportionately high levels of feeding on cats and dogs compared to humans. Richards et al. (2006) reported a similar trend for HFIs based on host abundance in North Carolina, but when time-weighted, found that humans were fed upon disproportionately often compared to cats and dogs^8^. In Brazil, HFIs based on host abundance showed the opposite trend to ours, suggesting that *Ae. albopictus* fed disproportionately often on humans compared to cats and dogs^13^. These results highlight the need for additional studies that measure host availability and also suggest a need for caution when extrapolating these results to make conclusions about innate mosquito preference. In both Long Island and North Carolina, collections were only conducted at a subset of houses per neighborhood, allowing for the movement of blood fed mosquitoes from properties where interviews were not conducted. Flight range for engorged blood fed *Ae. albopictus* is not known, but reported range of other blood fed species suggest that movement between properties is possible after feeding, as do records of *Ae. albopictus* dispersal between blood feeding and oviposition^39, 40, 41, 42^. Furthermore, household interview data depends on accurate self-reporting of outdoor activity, which may be unreliable^43^. This inaccuracy of outdoor time estimates is compounded if the interview is only administered once for the entire sampling period, such as in Richards et al. (2006)^8^.

We also assessed host availability through camera traps in order to calculate forage ratios for free-roaming animals, which suggest a tendency to feed on cats and opossums and to avoid raccoons, squirrels, and birds compared to their relative abundance in residential sites. While camera traps do not provide a perfect measure of host abundance, it is considered a robust method for mammal inventories^44^. Camera traps may be less useful in estimating bird abundance^45^, however, birds were one of the most frequently photographed groups of animals in our study, but were not fed upon, so improved accuracy in estimating bird abundance would not have altered conclusions drawn from forage ratio calculations.

Despite limitations, estimating host availability and abundance in conjunction with blood meal studies is much more informative than studies that lack such data. By understanding more about the context in which a certain feeding pattern arose, more general conclusions can be drawn about feeding behavior. The patterns revealed after accounting for host availability can be caused by numerous factors, such as host defenses. This may explain the high number of opossum blood meals because this nocturnal marsupial would likely be asleep, with decreased self-defense, during *Ae. albopictus* daytime biting activity. However, raccoons are also nocturnal and in contrast, were fed upon less often than expected, suggesting that innate preferences or other factors could potentially also be at play. Only two preference studies have been conducted for *Ae. albopictus*; in La Reunion Island, a no-choice blood feeding experiment on 12 animal species found chicken, human, dog and cow were fed upon more often than duck, shrew, rat, pig, mouse, goat, gecko, and chameleon^20^. Subsequently, a choice experiment showed higher attraction to humans compared to chicken, dog, cow and goat^20^. However, large and small animals were treated differently and were not given equal opportunities for self-defense, potentially affecting results. In Thailand, landing catches demonstrated preference for humans compared to pigs, buffalo, dogs, and chickens; however, the use of a second human to catch mosquitoes from the non-human animals may have impacted results. It therefore remains unclear whether *Ae. albopictus* has innate host preference.

One mechanism by which host preferences may evolve is through natural selection whereby feeding on a certain host enhances reproductive fitness, leading selection to favor genetic variants with preference for that host^46^. This is known to be the case for other species, such as *Ae. aegypti*^23^. We investigated the potential role of fitness in driving *Ae. albopictus* feeding patterns by assessing survival and egg production of mosquitoes fed on several host species in the Northeastern United States. Within the Long Island *Ae. albopictus* population, we found that host species had very limited impact on survival, egg production, or basic reproductive rate. The only significant differences were lower egg production after feeding on cats compared to humans and opossums, and no significant differences in survival. Interestingly, the reduced fecundity on cat blood is opposite to what we might expect based on the feeding index, which suggested a tendency to feed more often on cats compared to humans. A previous report from Baltimore of high feeding rates on rats, led us to compare the fitness of Long Island and Baltimore *Ae. albopictus* after feeding on human and rat blood. Specifically, we investigated whether differences in fitness may be driving the striking differences in feeding patterns between the two locations. However, the only significant difference was higher egg production by Long Island mosquitoes fed human blood than all three other groups. If egg production was driving this difference, we would expect to also see higher egg production for Baltimore mosquitoes fed rat compared to human blood, but this was not the case. Furthermore, survival of mosquitoes fed on human blood was longer than those fed on rat blood for both Baltimore and Long Island *Ae. albopictus*. Together, these results suggest that fitness advantage does not drive different feeding patterns in these two locations.

The impact of host species on *Ae. albopictus* egg production has only been assessed twice before. Gubler (1970) found greater fecundity for mouse-fed females, followed by guinea pig, rat, and chicken; however, the study was not replicated and no statistical analyses were conducted^25^. In another study, chicken-fed *Ae. albopictus* were less fecund than those offered guinea pig or human blood and, consistent with our results, no differences between the two mammal species were found^24^. These results do not demonstrate a selective pressure for *Ae. albopictus* to evolve preferences within mammalian hosts. However, preference can evolve through other pathways and should be assessed directly. Other specialist feeders lack apparent fitness advantages for their preferred host. For example, *Anopheles gambiae* has a well-established preference for humans, but in a single study conducted to date, there is no fitness advantage provided by a human-only diet compared to a generalist diet^47^.

It is also possible that when assessed under different conditions, differences in fitness by host species may be revealed. For instance, we did not provide the mosquitoes with sugar after blood feeding; the presence of sugar has been shown to reduce reproductive fitness in *Ae. albopictus* compared to human blood alone and mosquitoes on Long Island feed frequently on sugar^33, 48^. For *Ae. aegypti*, the addition of sugar changed the directionality of host species effects on fitness, shifting the fitness benefits from human to mouse blood^23^. If a similar phenomenon exists for *Ae. albopictus*, the absence of sugar in our experiments would maximize the fitness of human blood compared to other species. We also only provided the mosquitoes with one blood meal. Providing a more natural series of blood meals may have influenced our results.

*Aedes albopictus* is often referred to as anthropophilic due to the high percentage of human blood meals in numerous field studies and the preference assessments conducted by Delatte et al. (2010)^20^. However, this classification remains unproven. In fact, our results are more indicative of a generally mammalophilic feeding behavior for *Ae. albopictus*. It is important to understand the underlying blood feeding behavior and physiology of *Ae. Albopictus* because it influences and modulates the feeding patterns in the field, which will ultimately influence pathogen transmission^19^. In Long Island, the diverse utilization of hosts in residential and farm settings demonstrates that *Ae. albopictus* could serve as an enzootic bridge vector. However, the absence of bird blood meals suggests that *Ae. albopictus* may be of limited concern as a vector of West Nile and Eastern equine encephalitis viruses in the Northeastern US. Populations of *Ae. albopictus* in this region have sufficient vector competence to transmit numerous anthroponotic viruses^49, 50, 51^, but transmission of these pathogens may be limited due to lower rates of human feeding compared to other regions^52^.

Our results provide insight into disease transmission risk by *Ae. albopictus* in Northeastern United States. Additionally, our observations reveal that host availability has a major impact on feeding patterns, but did not fully explain blood meal distribution. Fitness benefits did not explain the feeding patterns observed in Long Island or Baltimore, highlighting the need for further research on determinants of *Ae. albopictus* feeding behavior.

## Acknowledgements

We thank Dr. Paul Leisnham at the University of Maryland and Dr. Shannon LaDeau at the Cary Institute for providing the Baltimore *Ae. albopictus* eggs, Dr. Alex Amaro for assistance with blood meal identification, Dr. Dan Gilrein at Cornell Cooperative Extension, Moses Cucura, and Dr. Scott Campbell at Suffolk County Government for providing laboratory space and logistical support, Dr. Erika Mudrak for providing statistical expertise, and Dr. Talya Shragai for assistance with field site selection. This material is based upon work that is supported by the National Institute of Food and Agriculture, U.S. Department of Agriculture, multistate Hatch project under 1014052. This work was also supported in part by Cooperative Agreement Number U01CK000509, funded by the Centers for Disease Control and Prevention. Its contents are solely the responsibility of the authors and do not necessarily represent the official views of the Centers for Disease Control and Prevention or the Department of Health and Human Services.

