## Supplemental Materials S1 for "The effects of host availability and fitness on *Aedes albopictus* blood feeding patterns in New York"

**Household interview**

1. Address:
2. How many people were living in this house over the past 2 days?
3. Were any animals staying here over the past 2 days?
4. (if yes) What type of animal? How many?
5. We are counting total human hours outside the house in the yard. How much time have people spent outside in your yard today, added all together?	(If prompting is needed, say “For example, if 2 people spent one hour outside so far today, that would be 2 hours”)
6. How much time did people spend outside in your yard yesterday?
7. How much time did people spend outside in your yard 2 days ago?
8. How much time did your pet(s) spend outside in your yard today?	Yesterday?	2 days ago?	*****Separate by type of animal; ie collect info separately for cats and dogs but add together time for 2 dogs.*
9. Did people wear mosquito repellent when they went outside over the past 2 days? (If yes, prompt “Always, usually or sometimes?”

Always	Usually	Sometimes	Rarely	Never

10. Did you do anything else to avoid mosquito bites, such as use insecticides during the last week?

Were windows open? Y / N	Did they have screens? Y / N

Were doors open? Y /	N	Did they have screen doors? Y / N
